# Human Herpesvirus 8 ORF57 protein is able to reduce TDP-43 pathology: Network analysis Identifies Interacting Pathways

**DOI:** 10.1101/2023.05.18.540717

**Authors:** Chelsea J Webber, Caroline N. Murphy, Alejandro N. Rondón-Ortiz, Sophie J.F. van der Spek, Elena X. Kelly, Noah M. Lampl, Giulio Chiesa, Ahmad S. Khalil, Andrew Emili, Benjamin Wolozin

## Abstract

Aggregation of TAR DNA-binding protein 43kDa (TDP-43) is thought to drive the pathophysiology of ALS and some Frontotemporal dementias. TDP-43 is normally a nuclear protein that in neurons translocates to the cytoplasm and forms insoluble aggregates upon activation of the integrated stress response (ISR). Viruses evolved to control the ISR. In the case of Herpesvirus 8, the protein ORF57 acts to bind protein kinase R, inhibit phosphorylation of eIF2α and reduce activation of the ISR. We hypothesized that ORF57 might also possess the ability to inhibit aggregation of TDP-43. ORF57 was expressed in the neuronal SH-SY5Y line and its effects on TDP-43 aggregation characterized. We report that ORF57 inhibits TDP-43 aggregation by 55% and elicits a 2.45-fold increase in the rate of dispersion of existing TDP-43 granules. These changes were associated with a 50% decrease in cell death. Proteomic studies were carried out to identify the protein interaction network of ORF57. We observed that ORF57 directly binds to TDP-43 as well as interacts with many components of the ISR, including elements of the proteostasis machinery known to reduce TDP-43 aggregation. We propose that viral proteins designed to inhibit a chronic ISR can be engineered to remove aggregated proteins and dampen a chronic ISR.

## Introduction

Tar DNA binding protein 43 (TDP-43) is a RNA binding protein (RBP) that accumulates in multiple neurodegenerative diseases, including amyotrophic lateral sclerosis (ALS), frontotemporal dementia-TDP-43 (FTD-TDP) and Alzheimer’s disease (AD) (1,2). TDP-43 normally resides in the nucleus but translocates to the cytoplasm during stress (3). Cytoplasmic TDP-43 interacts with other disease-linked RBPs, undergoes post-translational modifications and forms insoluble aggregates (4,5). Mutations in TDP-43 are sufficient to cause ALS, and mutations in other proteins linked to ALS and FTD-43 (e.g., C9orf72 or PGRN) are also known to lead to TDP-43 pathology. The causes of TDP-43 linked neurodegeneration appear to be complex. One major mechanism of TDP-43 induced degeneration result from the chronic stress arising from the prolonged accumulation of cytoplasmic TDP-43 and cytoplasmic TDP-43 aggregates. Another potential mechanism of degeneration arises from the loss of nuclear TDP-43, with a concomitant loss of nuclear splicing (6–12). Many therapeutic approaches focus on inhibiting the accumulation of cytoplasmic TDP-43 aggregates and inhibit nuclear to cytoplasmic translocation of TDP-43 (13–15), including by modulating phosphorylation of eIF2α (16,17).

Chronic stress elicits an integrated stress response (ISR), which inhibits RNA translation and catabolism. The ISR is beneficial in addressing acute stresses, but in human disease a chronic, persistent stress response can itself contribute to the disease process (18). In neurodegenerative disease chronic activation of the ISR leads to prolonged accumulation of RNA-protein complexes, termed stress granules (SGs). These SGs act as a conduit in a pathway that produces cytoplasmic TDP-43 aggregates. The ISR pathway can be activated by four kinases that respond to exogenous and endogenous stressors: general control nonderepressible 2 (GCN2) kinase (19), heme-regulated eIF2α kinase (HRI) (20), PKR-like endoplasmic reticulum kinase (PERK) (21), and protein kinase R (PKR) (22). Both PERK and PKR have been shown to play important roles in the pathophysiology of neurodegenerative diseases, with inhibitors of these enzymes able to reduce the pathophysiology in many models of neurodegenerative diseases, including models of ALS and frontotemporal dementia (FTD), which express TDP-43 (23–25). Despite this promise, inhibitors of PERK and PKR have failed in the clinic because of peripheral toxicity (26,27).

An alternative approach to modulating the ISR for therapeutic purposes might be to take advantage of proteins naturally evolved to modulate proteostasis or inhibit the ISR. Prior studies have examined the plant Arabidopsis protein HSP104, which acts as a powerful disaggregase of disease linked proteins (28–31). However, HSP104 has not been adapted by nature to regulate the mammalian proteostasis machinery. In contrast, viral proteins evolved to target many elements of the mammalian ISR in order to maintain protein synthesis and allow production of viral proteins. For instance, Herpesvirus 8 produces the protein ORF57, which inhibits PKR activity and the ISR by binding and inhibiting both PKR and PKR activating protein, and is adapted to function in neurons (32–35). By inhibiting the ISR, ORF57 also allows a neuron to maintain normal translation, splicing and RNA stability, despite the impact of the virus on a cell (36–38). We hypothesized that, although the ability of ORF57 to inhibit the ISR is problematic for our bodies in the case of a viral infection, OSR57 expression might be beneficial in the case of a chronic neurodegenerative disease.

In this study we used a genetically engineered SH-SY5Y neuroblastoma cell line to model the actions of TDP-43 in neurodegenerative diseases. Two SH-SY5Y cell line were engineered: one constitutively expresses wild type TDP-43, while the other expresses a tetracycline inducible TDP-43 construct lacking its nuclear localization signal. Of note, the inducible TDP-43 lines, termed TDP-43ΔNLS, readily forms cytoplasmic aggregates in response to induction plus stress, modeling what occurs in neurons of patients with ALS, FTD-TDP or AD. We investigated the effects of ORF57 on aggregate formation and dispersal in the TDP-43ΔNLS SH-SY5Y line. We demonstrate that ORF57 interacts with proteins robustly involved in TDP-43 aggregation, promotes the removal of TDP-43 aggregates and reduces TDP-43-mediated neurodegeneration. These results suggest that ORF57 might be able to be utilized as a therapeutics for neurodegenerative diseases exhibiting TDP-43 pathology.

## Results

### ORF57 is protective against oxidative stress in human neuroblastoma cells exhibiting TDP-43 pathology

TDP-43 is a particularly prominent RBP that is linked to multiple neurodegenerative diseases, including ALS, FTD-TDP and AD. While predominantly a nuclear RBP, TDP-43 aggregates in the cytoplasm in these diseases. We model the disease related cytoplasmic TDP-43 aggregates by overexpressing a TDP-43 construct lacking a functional nuclear localization signal, termed TDP-43ΔNLS (39). In cell culture, this construct yields abundant cytoplasmic TDP-43 which aggregates in response to activation of the ISR with sodium arsenite (**SA**) treatment (39). ORF57 inhibits the ISR through known interactions with PKR (32).

We proceeded to examine how ORF57 affects the ISR in SH-SY5Y cells with inducible expression of TDP-43ΔNLS. The SH-SY5Y cells were transduced with ORF57::mCherry or mCherry expressing lentiviruses and the transgene expression level was quantified by mRNA levels (**Fig. 1A, B**). After infection, TDP-43ΔNLS was induced by treatment with doxycycline (1 μM, 24 hrs), and then the SH-SY5Y cells expressing TDP-43ΔNLS ± ORF57 were subjected to SA stress (300 μM, 90 min). SA stress activates PKR to phosphorylate eIF2α (**p-eIF2**α) and halts global translation. We confirmed that these effects of SA stress were relieved by ORF57 expression. The TDP-43ΔNLS SH-SY5Y cells exhibited increased levels of p-eIF2α in response to SA stress (**Fig. 1C, D**), this was also observed in wildtype SH-SY5Y cells (**Supp. Fig. 1A, B**). Treatment with SA also induced abundant cytoplasmic TDP-43 granules (**Fig. 2A, F, G**). We found that ORF57 expression relieved multiple ISR associated phenotypes. ORF57 expression reduced p-eIF2α by 45% (P<0.0001) in SH-SY5Y TDP-43ΔNLS cells following exposure to SA (**Fig. 1C, D**). We found that ORF57 increased protein synthesis when measured with the SUnSET assay (P<0.05) (**Fig. 1E, F**), however this increase was small when compared to non-stressed TDP-43 overexpressing cells (**Supp. Fig. 2**). This marginal increase in protein synthesis was also observed by SUnSET assay in wildtype SH-SY5Y cells (**Supp. Fig. 1C-F**). ORF57 also protected against apoptosis in SH-SY5Y and TDP-43 NLS cells (**Fig. 1G-J**). Cleaved caspase-3 levels were significantly reduced in ORF57 compared to mCherry expressing cells in both stress and basal conditions (P<0.05), demonstrating that ORF57 is protective against TDP-43 NLS induced apoptosis. Similar effects were observed in wildtype SH-SY5Y cells expressing ORF57 (**Fig. 1I, J**).

**Figure 1.**
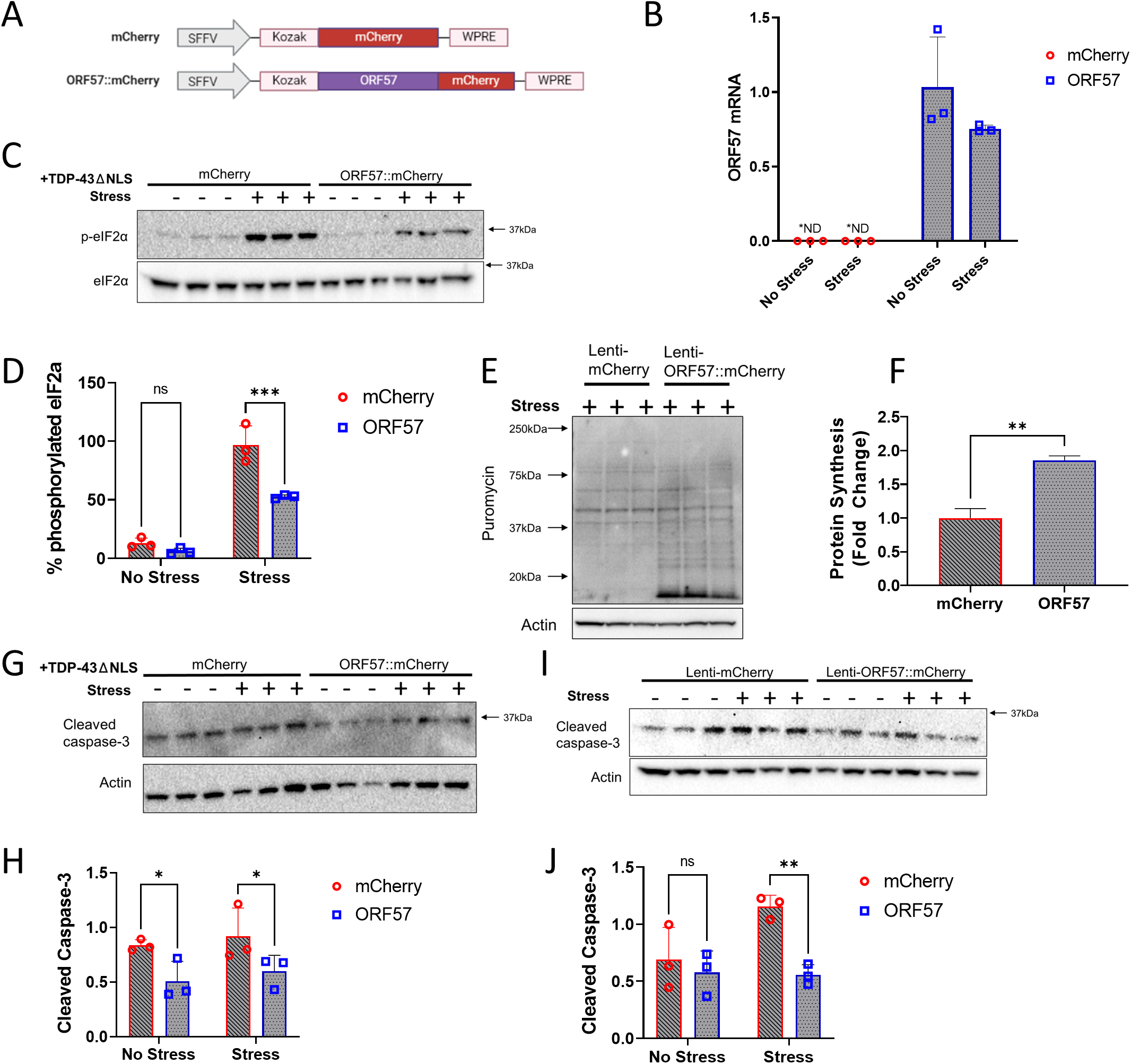
ORF57 protects against oxidative stress in TDP-43ΔNLS overexpressing SH-SY5Y cells. Experiments conducted in mCherry and ORF57 mCherry expressing SH-SY5Y TDP-43ΔNLS cells with and without SA stress. **A)** Constructs for mCherry and ORF57::mCherry lentivirus. **B)** Overexpression of ORF57 in SH-SY5Y cells by rt-qPCR. **C)** Western blot for phosphorylated eIF2α and total eIF2α. **D)** Quantification of C. **E)** SUnSET of mCherry and ORF57 expressing TDP-43ΔNLS cells with SA stress. **F)** Quantification of E. **G)** Western blot of cleaved caspase-3 in TDP-43ΔNLS cells. **H)** Quantification of G. **I)** Western blot of cleaved caspase-3 in non-overexpressing SH-SY5Y cells. **J)** Quantification of I.

**Figure 2.**
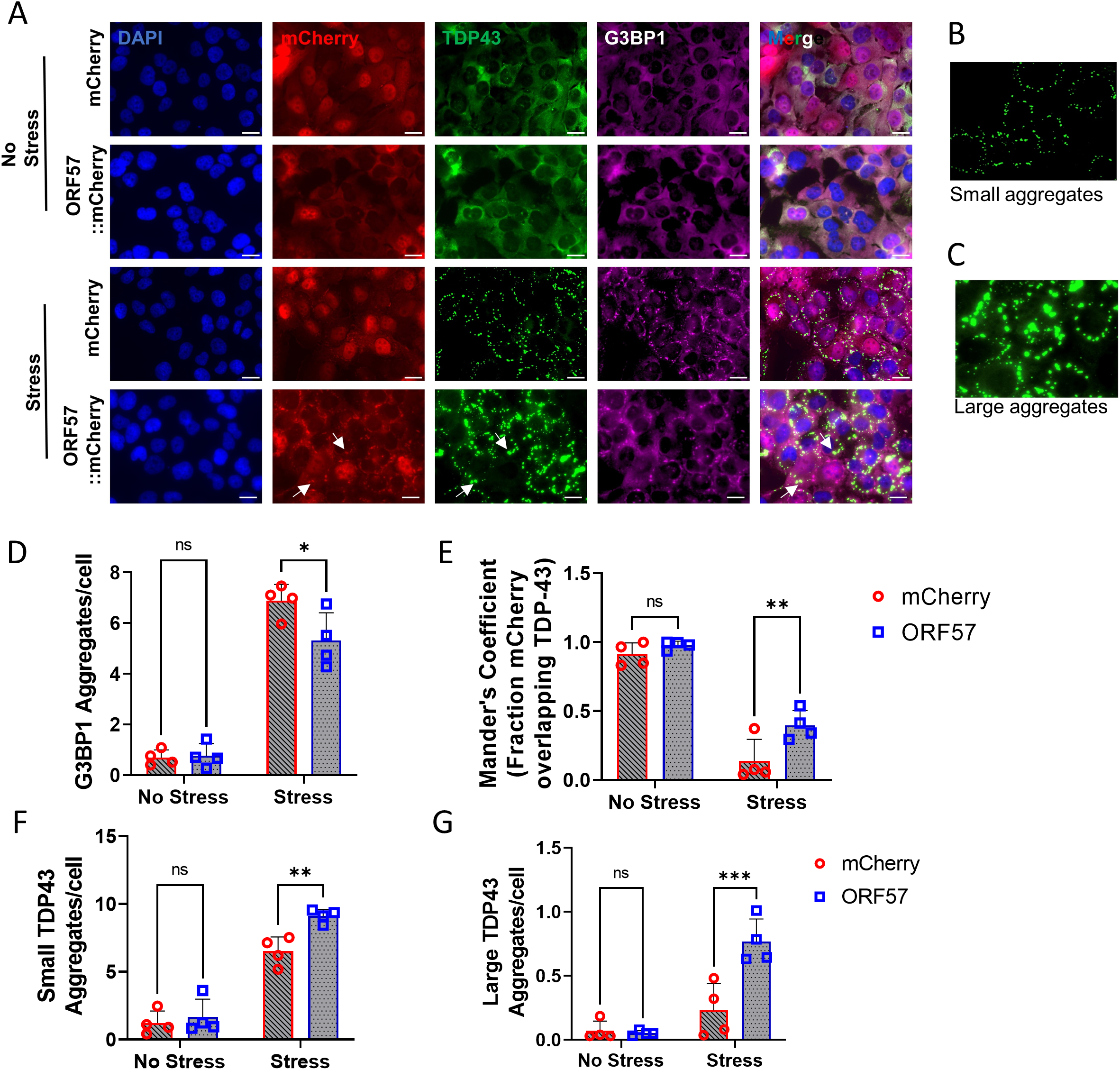
Stress granule assembly and TDP-43 aggregation dysregulated by ORF57. **A)** Immunocytocemistry in SH-SY5Y TDP-43ΔNLS. DAPI (405 nm). mCherry or ORF57::mCherry (594 nm). TDP-43ΔNLS::GFP (488 nm). G3BP1 (647 nm). Merge, overlay of all images. White arrows indicate overlap between ORF57 and TDP-43 aggregates. **B)** Representation of small TDP-43ΔNLS aggregates 10-200 Aggrecount units. **C)** Representation of large TDP-43ΔNLS aggregates 200-400 Aggrecount units. **D)** Quantification of G3BP1 aggregates in each cell condition from A. **E)** Mander’s coefficient of fraction of mCherry overlapping with TDP-43 signal. **F)** Quantification of small TDP-43ΔNLS aggregates. **G)** Quantification of large TDP-43ΔNLS aggregates. Two-way ANOVA with Bonferroni post hoc comparison, *P<0.05, **P<0.01, ***P< 0.0001.

SG formation was also reduced with ORF57 expression. Imaging was performed on induced SH-SY5Y TDP-43ΔNLS cells expressing ORF57::mCherry or mCherry alone. G3BP1 positive SGs were reduced >20% (P<0.05) in cells expressing ORF57 compared to those expressing mCherry under SA stress (**Fig. 2A, D**). As described above SA stress also caused cytoplasmic TDP-43 to aggregate (**Fig. 2A–C, F, G**). We observed that ORF57 appeared to be colocalizing with TDP-43 (**Fig 2A, white arrows**) and therefore we analyzed colocalization of mCherry with TDP-43 signals. The fraction of the mCherry signal in both mCherry and ORF57::mCherry that overlapped with the unstressed TDP-43 signal was more than 0.9 (**Fig. 2E)**. However, once stress was added the fraction of the mCherry signal that overlapped with TDP-43 was dependent on ORF57 expression. Stress caused the TDP-43 to form cytoplasmic inclusions without altering the mCherry signal, therefore the fraction of mCherry overlapping with TDP-43 signal was reduced to 0.137 ± 0.158. Meanwhile, the ORF57::mCherry signal was changed in response to stress and the fraction of ORF57::mCherry overlapping with the TDP-43 signal was 0.396 ± 0.106 (P<0.05). This indicates that ORF57 colocalizes with aggregated TDP-43.

Co-expressing ORF57 overall increased the number of TDP-43 aggregates and changed the structure of the TDP-43 aggregate. ORF57 caused TDP-43 to form large amorphous aggregates that were almost completely absent in mCherry control cells (**Fig. 2A–C, F, G**). Parameters were made to distinguish these aggregates for quantification, marking the small aggregates from 10-200 Aggrecount units and the large aggregates 200-400 units. According to these parameters, small and large aggregates were quantified. Under stress, small aggregates of TDP-43 were increased by approximately 30% (P<0.05) in ORF57 expressing cells compared to mCherry expressing cells. Large aggregates of TDP-43 were also increased in ORF57 cells under stress compared to mCherry expressing cells (P<0.05).

### ORF57 reduces TDP-43 aggregation

We hypothesized that the large TDP-43 aggregates represent a more diffuse state, and therefore contain less insoluble and more soluble TDP-43. In order to assess this, we fractionated TDP-43 into RIPA-soluble and RIPA-insoluble fractions. Expression of ORF57 improved TDP-43 solubility during stress, increasing the amount of soluble and decreasing the amount of insoluble TDP-43 (**Fig. 3A-C**). Under basal conditions, approximately 87% of TDP-43 was in the soluble fraction in both ORF57 and mCherry treated cells (**Supp. Fig. 3**). With SA stress, the amount of soluble TDP-43 in mCherry cells was reduced to 59% and the amount of insoluble TDP-43 was increased 41%. ORF57 expression prevented the TDP-43 from becoming insoluble. In ORF57 cells, 75% of TDP-43 was in the soluble fraction and only 26% was insoluble P<0.05) (**Fig. 3A-C**). These results indicate that ORF57 inhibits transition of TDP-43 to the insoluble fraction, which occurs in response to SA stress.

**Figure 3.**
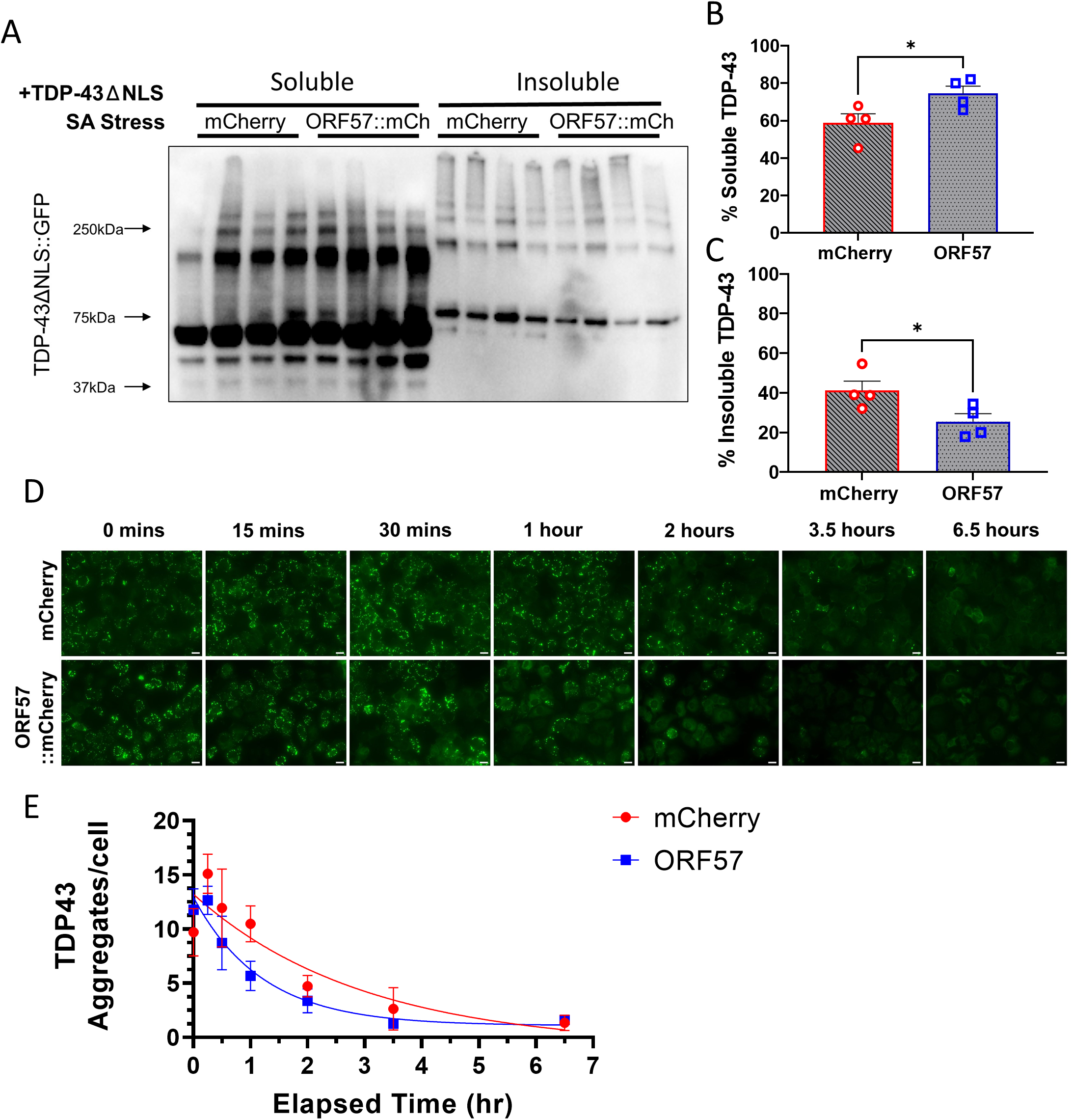
TDP-43 aggregates are more soluble with ORF57. Experiments conducted in mCherry and ORF57 mCherry expressing SH-SY5Y TDP-43ΔNLS cells with SA stress. **A)** Western blot of TDP-43ΔNLS::GFP RIPA-soluble and in-soluble fractions. **B)** Quantification of %RIPA-Soluble TDP-43ΔNLS::GFP calculated as soluble TDP-43 divided by the sum of soluble and insoluble TDP-43. **C)** Quantification of %RIPA-Insoluble TDP-43ΔNLS::GFP. **D)** Time course experiment measuring TDP-43ΔNLS::GFP aggregate recovery at 15 minutes, 30 minutes, 1 hour, 2 hours, 3.5 hours, and 6.5 hours after SA removal. **E)** Quantification of E with exponential one-phase decay best fit curves. Unpaired t-tests, *P<0.05. Scale bars 20µm.

Next, we hypothesized that the ability of ORF57 to increase soluble and reduce insoluble TDP-43 might lead to more rapid resolution of TDP-43 granules upon removal of stress. To investigate this, we conducted a time course experiment to quantify the kinetics of TDP-43 granule resolution after SA stress. SH-SY5Y TDP-43 NLS cells expressing ORF57 or mCherry were subjected to SA stress (300 M, 90 min), and then SA was removed. The total number of TDP-43 aggregates/cell were quantified at 15 minutes, 30 minutes, 1 hour, 2 hours, 3.5 hours, and 6.5 hours after removal of SA stress. The SH-SY5Y TDP-43 NLS cells expressing ORF57 exhibited a faster resolution of TDP-43 granules than mCherry expressing cells (**Fig. 3D, E**). The T1/2 for recovery was 0.84 hr for the ORF57 cells vs 2.06 hr for the mCherry cells, which is a 2.45 fold (P<0.017) difference using a one-phase decay nonlinear regression model (**Fig. 3D, E**). This data indicates that ORF57 protects against insoluble TDP-43 aggregation in this model.

### ORF57 binds to stress pathway proteins involved in TDP-43 pathogenesis

In order to gain insight into the mechanisms through which ORF57 might affect TDP-43 granule dynamics, we investigated the ORF57 interactome in SH-SY5Y cells expressing ORF57 under basal and stressed conditions. Prior literature shows that ORF57 binds to PKR (Gene ID: *EIF2AK2*), but does not explore whether ORF57 might also bind other proteins linked to protein aggregation.

Using the mCherry-tagged TDP-43ΔNLS SH-SY5Y cell line, we were able to explore the ORF57 protein interaction with proteins involved in the ISR as well as those involved in TDP-43 aggregation. The cells were transduced with mCherry or ORF57::mCherry and immunoprecipitated with anti-RFP nanobody. Following washing and elution, the resulting eluant was analyzed by mass spectrometry (IP-MS) under basal overexpression of cytoplasmic TDP-43 conditions as well as SA-induced cellular stress (**Fig. 4A**). Under basal conditions, IP-MS revealed binding of ORF57 with 196 proteins (FDR-adjusted P-value < 0.05 and log2 fold-change > 0.58 versus the mCherry control) (**Fig. 4B**). Importantly, these interactions included PKR, encoded by the *EIF2AK2* gene, which is a known ORF57 interactor and initiator of the ISR (**Fig. 4B**). Enriched Gene Ontology (GO) Biological Process (BP) functional annotation terms revealed basal conditions binding of ORF57 to proteins involved in Stress Response (SRP) –dependent co-translational protein targeting to membrane, mRNA metabolism, RNA translation and RNA processing (**Fig. 4B**). These functional terms are consistent with known functions of ORF57 as a regulator of viral protein synthesis. Under SA-induced cellular stress, ORF57 revealed binding with 144 proteins (FDR-adjusted P-value < 0.05 and log2 fold-change > 0.58 versus mCherry-SA control) (**Fig. 4C**; **Supp. Table 1**). There was high overlap of ORF57 bound-proteins between basal and SA conditions (133 proteins, Venn diagram) (**Fig. 5A**). There were 28 proteins that interacted with ORF57 greater than 4-fold either under TDP-43 overexpression or with the addition of stress (**Table 2**). Expectedly many of these proteins are involved in the antiviral response: TRIM25, ZC3HAV1, PABPC4, HELZ2, SYNCRIP, DHX30, YBX3, EIF2AK2, RBMX, and DHX37. While other proteins are involved in SG assembly and disease pathology including HNRNPA2B1, FUS, TAF15, HNRNPDL, PABPC1, DHX30, ELAVL1, MTDH, HNRNPA3, EIF2AK2, HNRNPR, and MATR3. These high interactors help to indicate the primary function of ORF57 under conditions of TDP-43 pathology.

**Figure 4.**
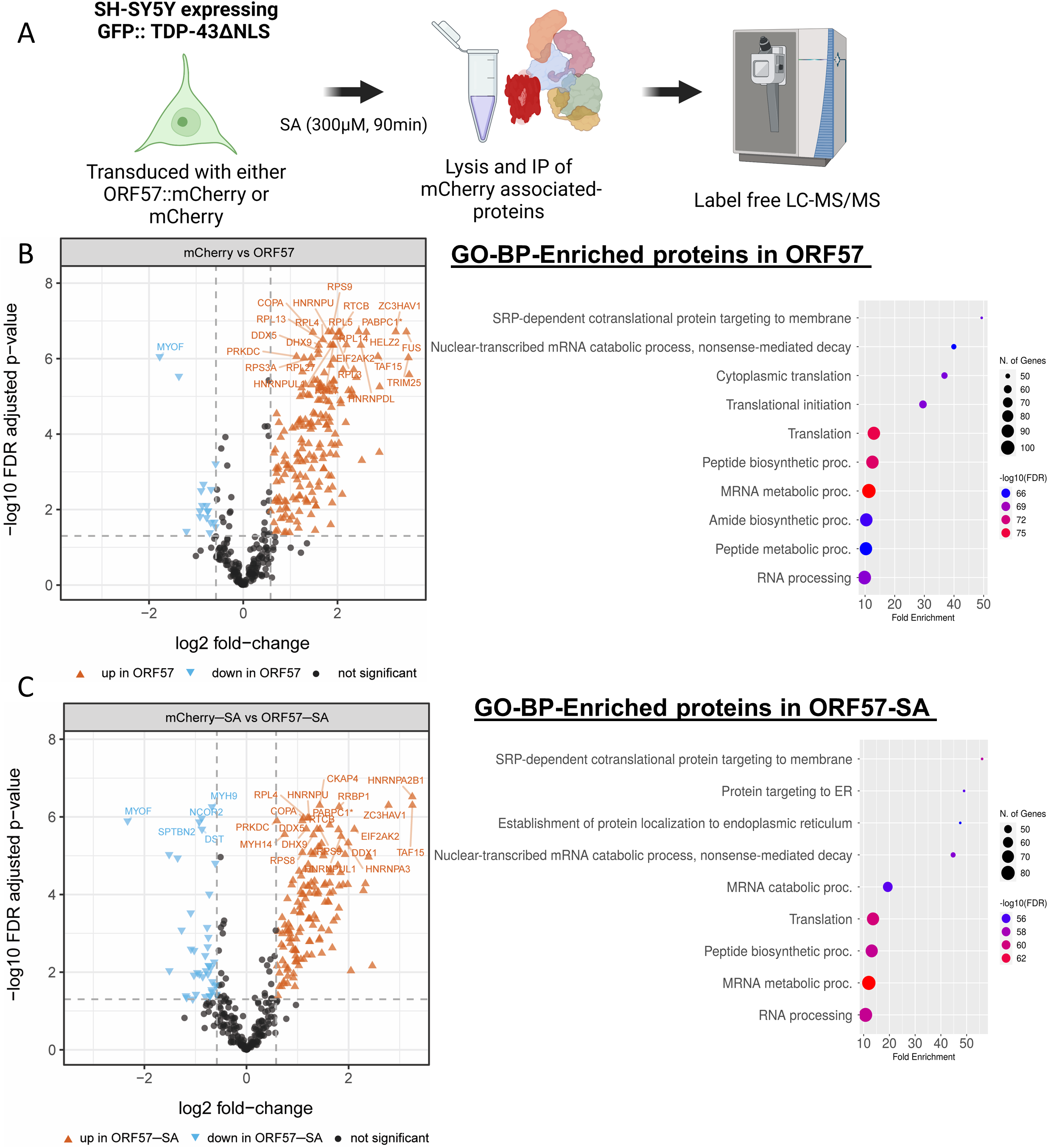
ORF57 interactome. **A**) Outline of proteomic approach using SH-SY5Y cells expressing GFP::TDP-43ΔNLS. **B)** High-confidence ORF57-bound proteins observed under basal conditions and the GO-BP functional annotations terms associated with these enriched proteins. Functional annotation terms were determined using ShinyGO Version 0.76.3 (77) **C)** High-confidence ORF57-bound proteins observed in presence of SA and the GO-BP functional annotations terms associated with these enriched proteins. Enriched proteins in basal and stress conditions were considered significant when they have a Log2 fold-change higher than 0.58 and an FDR adjusted P-value less than 0.05.

**Figure 5.**
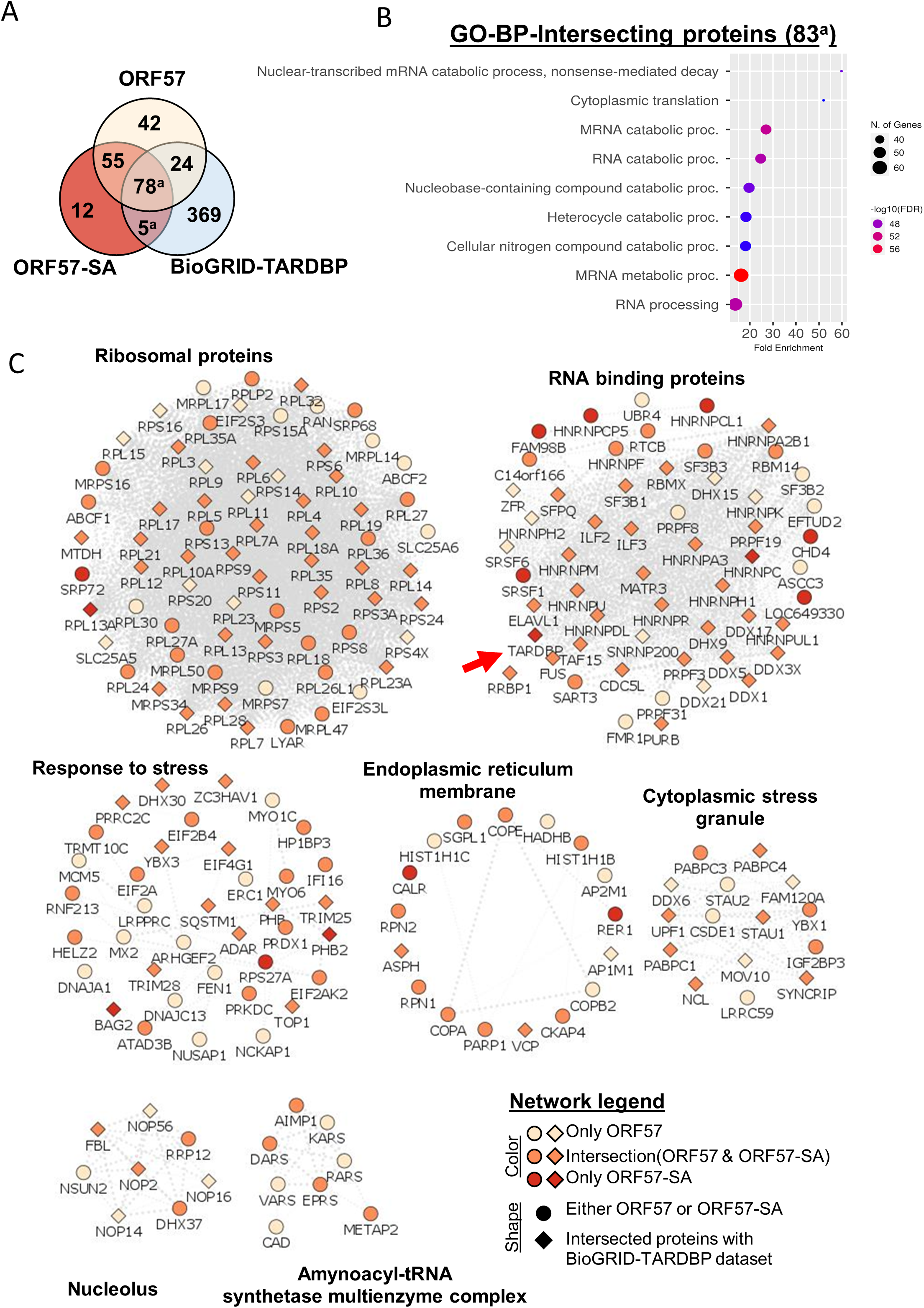
ORF57 interacts with theTDP-43 protein network. **A)** Venn diagram using ORF57, ORF57-SA and the TDP-43 BioGRID-TARDBP) datasets. **B)** Functional annotations terms for overlapping proteins between ORF57-SA and BioGRID-TARDBP datasets. **C)** Multi-module representation of ORF57-bound protein networks (Using Cytoscape, STRING-db and ClusterMaker2. Node title describes gene names, node shape describes whether the protein was found in either ORF57 datasets (circle) or BioGRID-TARDBP (diamond), and color indicates whether the gene was only found in either ORF57, ORF57-SA or overlapped between them.

**Table 1.**
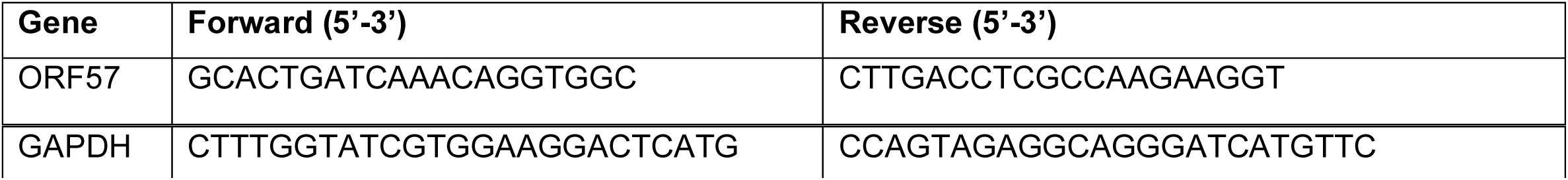
rt-qPCR primers for ORF57.

**Table 2.** Proteins that interact highly with ORF57.

more than a 4-fold increase in binding either in the absence or presence of sodium arsenite (SA) stress. All proteins have FDR-adjusted p-value <0.05.
| **Protein** | **Log-2 Fold Change**<br>**mCherry vs ORF57** | **Log-2 Fold Change**<br>**mCherry-SA vs ORF57-SA** |
| --- | --- | --- |
| HNRNPA2B1 | 3.521 | 3.248 |
| TRIM25 | 3.506 | 1.640 |
| FUS | 3.462 | 2.391 |
| ZC3HAV1 | 3.235 | 2.782 |
| RTRAF | 2.894 | 2.329 |
| PABPC4 | 2.888 | 1.609 |
| TAF15 | 2.859 | 3.249 |
| HELZ2 | 2.611 | 2.291 |
| SYNCRIP | 2.517 | 1.693 |
| HNRNPDL | 2.497 | 1.643 |
| PABPC1;<br>PABPC3 | 2.450 | 1.630 |
| RPL19 | 2.401 | 1.397 |
| DHX30 | 2.345 | 2.465 |
| ELAVL1 | 2.336 | 1.337 |
| MTDH | 2.297 | 1.816 |
| YBX3 | 2.286 | 0.802 |
| HNRNPA3 | 2.234 | 1.803 |
| EIF2AK2 | 2.189 | 2.113 |
| RRP12 | 2.175 | 1.633 |
| ASPH | 2.117 | 1.630 |
| RBMX | 2.101 | 0.962 |
| RPL14 | 2.081 | 0.983 |
| HNRNPR | 2.074 | 1.064 |
| DHX37 | 2.062 | 1.791 |
| RTCB | 2.046 | 1.384 |
| RPL5 | 2.023 | 1.471 |
| MATR3 | 1.887 | 2.045 |

The addition of SA stress also changed many proteins binding to ORF57. There were 4 proteins that significantly changed their ORF57 interaction in the presence of SA stress that were unchanged in mCherry. Three proteins (PRDX1, TRIM28, EIF2S3) significantly increased their ORF57 binding while 1 protein (NUSAP1) significantly decreased its binding with ORF57 with stress (**Table 3**).

**Table 3.** Unique proteins whose ORF57 interactions are significantly altered in the presence of stress.

Proteins that significantly change their interaction with ORF57 in the presence of sodium arsenite (SA) stress. These proteins are not significantly changed in mCherry vs mCherry-SA comparisons.
| **Protein** | **Log-2 Fold Change**<br>**ORF57 vs ORF57-SA** | **FDR-adjusted p-values** |
| --- | --- | --- |
| PRDX1 | 1.494 | 1.25E-06 |
| TRIM28 | 0.771 | 2.73E-03 |
| EIF2S3;<br>EIF2S3L;<br>F8W810 | 0.635 | 1.27E-03 |
| NUSAP1 | -1.169 | 1.18E-03 |

The ORF57 interactome network showed 17 proteins that bind to ORF57 only under stress conditions (**Table 4**). The 17 proteins have diverse functions within the cell which highlights the ability of ORF57 to target many different biological processes. The ORF57 interactome network was visualized using Cytoscape and String-db; this network was then sub-clustered to create multiple modules (**Fig. 5C**). Importantly, the network of proteins specific to the ORF57-SA interactome includes TDP-43 (gene name *TARDBP)* (**Fig. 5C, red arrow**). The ORF57-SA specific interactome network also includes other proteins known to be implicated in the ISR such as, CALR, RER1, and RBPs such as HNRNPC & CL1, HRNPCP5, SRSF1, RPL13A, RPS27A (**Fig. 5C**). The ORF57 interactomes show unique GO BP functional annotation terms from ORF57-SA that were not observed under basal conditions including: protein targeting to endoplasmic reticulum (ER) and mRNA catabolic processes. Proteins such as CALR and RER1 are ER proteins that explain the ER-associated functional GO terms displayed in Fig. 4C. The module “Response to stress” contains many proteins that are associated with proteostasis (*e.g.,* SQSTM1, DNAJ13, TRIM28, RPS27A, and others) and the ISR (*e.g.* EIF2AK2, EIF2A, etc.). This module could contribute strongly to the ability of ORF57 to inhibit the ISR and reduce TDP-43 aggregation. There have been at least 42 genes that have mutations which cause ALS disease (40). In our proteomics study, we found ORF57 to interact with 7 of these ALS implicated proteins in our TDP-43 pathology cells (**Table 5**). Sodium arsenite stress significantly increased ORF57’s interaction with TARDBP but not with the other 6 ALS proteins. Stress also significantly increased TARDBP’s interaction with mCherry. Overall, ORF57 interacted with 196 and 144 proteins for basal and stress conditions, respectively (**Supp. Table 1**).

**Table 4.** Proteins that are unique interactors of ORF57 in the presence of stress.

Proteins that interact significantly more in the presence of sodium arsenite (SA) stress with ORF57 than mCherry.
| **Protein** | **Log-2 Fold Change**<br>**compared to mCherry-SA** | **FDR-adjusted p-values** |
| --- | --- | --- |
| HNRNPC;<br>HNRNPCL1;<br>HNRNPCL2;<br>HNRNPCL3;<br>HNRNPCL4 | 0.616 | 0.04018 |
| CALR | 0.688 | 0.02346 |
| BAG2 | 0.702 | 0.00102 |
| RER1 | 0.807 | 0.01270 |
| RPS27A | 0.959 | 0.00530 |
| SRSF1 | 0.961 | 0.00267 |
| PHB2 | 0.974 | 0.00139 |
| DNAAF5 | 0.708 | 0.00507 |
| TARDBP | 0.721 | 0.00227 |
| SRP72 | 1.244 | 0.00043 |
| CHD4 | 1.328 | 0.00035 |
| FAM98B | 1.809 | 0.00022 |
| RPL13A | 1.818 | 0.00001 |

**Table 5.**
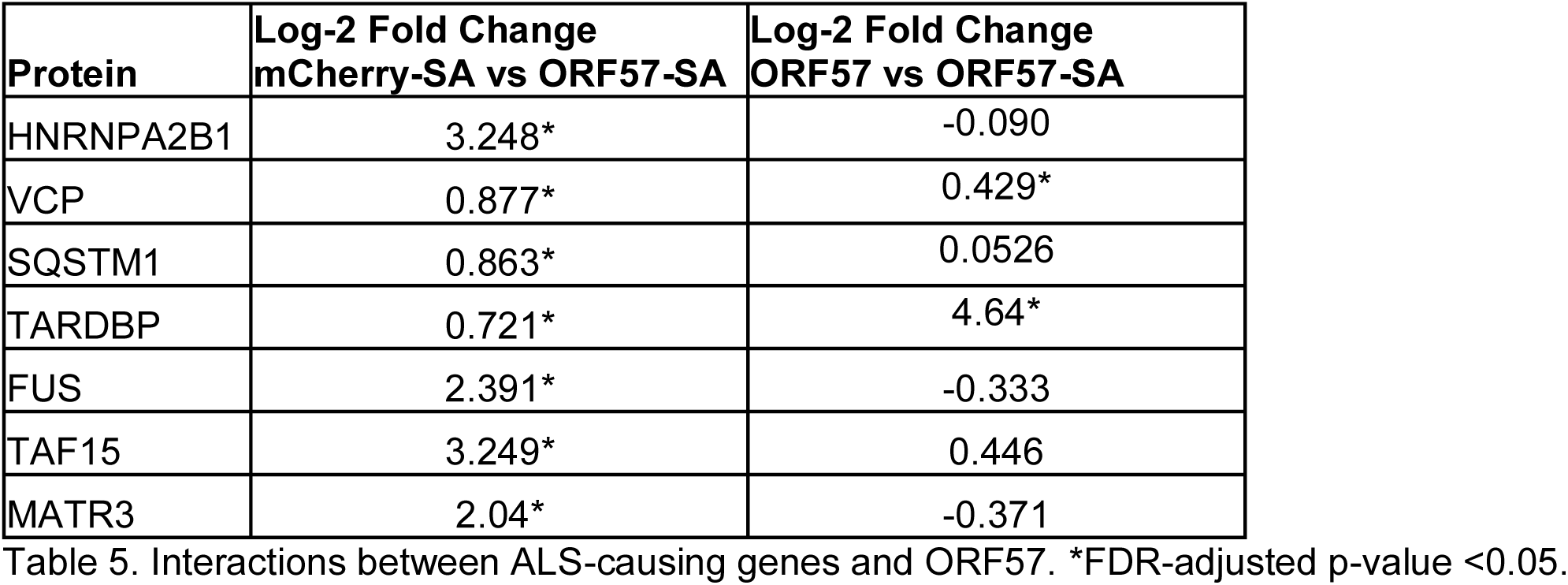
Amyotrophic lateral sclerosis disease linked proteins that interact significantly with ORF57.

Interactions between ALS-causing genes and ORF57. \*FDR-adjusted p-value <0.05.
| **Protein** | **Log-2 Fold Change**<br>**mCherry-SA vs ORF57-SA** | **Log-2 Fold Change**<br>**ORF57 vs ORF57-SA** |
| --- | --- | --- |
| HNRNPA2B1 | 3.248* | -0.090 |
| VCP | 0.877* | 0.429* |
| SQSTM1 | 0.863* | 0.0526 |
| TARDBP | 0.721* | 4.64* |
| FUS | 2.391* | -0.333 |
| TAF15 | 3.249* | 0.446 |
| MATR3 | 2.04* | -0.371 |

## Discussion

Aggregation of TDP-43 plays important roles in the pathophysiology of multiple diseases, including ALS, FTD-TDP and AD. The experiments described above used neuronal SH-SY5Y cells to demonstrate that ORF57 binds to TDP-43, proteins linked to TDP-43 proteostasis, proteins involved in the ISR, translation proteins, and PKR. We further demonstrated that expressing ORF57 elicited a striking 2.45-fold increase in the rate of TDP-43 granule dispersion and a 40% reduction in levels of cleaved caspase 3, which suggest that ORF57 is neuroprotective. We also observed increased protein synthesis in the presence of induced TDP-43 and changes in the morphology of TDP-43 cytoplasmic granules. Taken together these data suggest that TDP-43 might be able to act as an agent that can disperse TDP-43 granules and inhibit TDP-43-mediated neurodegeneration.

The studies of the ORF57 protein interactome network provide strong insights into the mechanisms through which ORF57 might impact on TDP-43 aggregation. ORF57 was previously shown to inhibit PKR and reduce SG accumulation, but these studies were not done in the context of TDP-43 aggregation. Multiple studies suggest that TDP-43 first consolidates in SGs, and then subsequently proceeds to aggregate (5,41,42). Our proteomic studies of ORF57 show that this protein binds to TDP-43, as well as other known binders of TDP-43, and regulate TDP-43 proteostasis (43). Some of these proteins, such as SQSTM1 and VCP, are directly involved with clearance of TDP-43 through autophagy and these proteins exhibit mutations associated with TDP-43-opathies (44–52). Other proteins are chaperones with well documented roles in proteostasis, such as DNAJA1, DNAJC13 and BAG2, and the ubiquitin transferase TRIM25 (53–56). Our proteomics confirm binding of ORF57 to known interactors SRSF1 (57) and PABPC1 (58,59), which are RBPs also known to associate with TDP-43 and regulate translation (60,61). Binding to stress proteins is readily apparent in the ORF57 interactome sub-module titled “Response to stress”, which includes many classic stress linked proteins including EIF2A, EIF2AK2, EIF4G1 and EIF2B4 (**Fig. 5C**). ORF57 also showed a strong ability to inhibit phosphorylation of EIF2, which is consistent with prior studies showing that it binds PKR and inhibits PKR activity (32–35). The presence in the ORF57 network of other RBPs that are known risk factors for neurodegenerative diseases, including FUS, HNRNPA2B1, MATR3 and TAF15 raises the possibility that the beneficial actions of ORF57 might extend beyond TDP-43 (62–66). These many interactions point to broad mechanisms through which ORF57 could impact to reduce aggregation of TDP-43 and promote granule resolution.

The clearest actions of ORF57 lie in the resolution of TDP-43 granules and the reduction in cleaved caspase 3, suggesting that ORF57-mediated reduction of TDP-43 aggregation is associated with a reduction in cell death. The protein interaction network for ORF57 shows that it interacts with many proteins known to regulate SG formation. However, this interactome impacts on SGs and translational control in a manner that appears to be complex. For instance, although ORF57 increased removal of TDP-43 aggregates, it actually increased the number of granules, with about a 3.5-fold increase in the number of large, diffuse looking granules (**Fig. 2G, J**). The diffuse impression of the SGs was consistent with solubility studies of TDP-43 showing that ORF57 increased the ratio of soluble to insoluble TDP-43 by about 50% (**Fig. 3A, B**). One possibility is that ORF57 stabilized soluble TDP-43 in SGs, reducing their exit to form insoluble TDP-43 aggregates.

Multiple studies have investigated putative links between infection with Herpes simplex virus type I (HSV-1) and neurodegenerative diseases, including Alzheimer’s disease and ALS. Relevant pathological studies suggest that HSV-1 can increase production of β-amyloid, tau phosphorylation and local inflammation (67–70); however HSV-1 infection has not been observed to induce intracellular TDP-43 or tau aggregates (40,67,69,71,72). Selective transduction of ORF57 differs studies of intact viruses because we used one particular gene, which was specifically selected for its potential to reduce TDP-43 aggregation. The work presented in this study demonstrates that proteins produced by Herpes viruses, such as ORF57, have the ability to limit the accumulation of protein aggregates through mechanisms shown above.

A limitation of this study is the use of immortalized cell lines over-expressing TDP-43. The approach of over-expressing ΔNLS TDP-43 is required when using neuroblastoma lines because TDP-43 does not form cytoplasmic aggregates under conditions of endogenous expression. Future studies will need to examine the actions of ORF57 in neuronally differentiated iPSC lines from patients who had sporadic or familial ALS, which do appear to exhibit TDP-43 cytoplasmic translocation and granule formation spontaneously or in response to stress.

Viruses evolved to control the translational machinery in order to induce the host cells to produce their viriomes. ORF57 provides one example of a protein that functions explicitly in regulating the host translational machinery, but there are many other proteins in other viruses that serve the same function. Our studies suggest that these pathways share strong overlap with pathways involved in protein aggregation processes that occur in neurodegenerative diseases. The work presented in this manuscript shows how viral proteins might also be engineered to yield new therapeutic approaches for neurodegenerative disease.

## Methods

### Cell Culture

Two human neuroblastoma, SH-SY5Y, cell lines were used for this study: regular SH-SY5Y and SH-SY5Y with a doxycycline-inducible TDP43 with its nuclear localization signal mutated (TDP-43ΔNLS). The TDP-43ΔNLS was a gift provided by Glenn Larsen, Ph.D., Aquinnah Pharmaceutical Inc.; requests for use of the cell line will be made at the determination of Aquinnah Pharmaceuticals, Inc.. Both cell lines were maintained in complete media containing Dulbecco’s Modified Eagle Medium with high glucose (Corning 10-013-CV), 10% tet-tested fetal bovine serum, 1% penicillin streptomycin (Gibco Pen Strep, # 15070-063), 1% non-essential amino acids (Gibco MEM NEAA, # 11140-050), and 1% Glutamine. TDP-43ΔNLS overexpression was induced with 1 µg/mL doxycycline added to complete media for 24 hours.

### ORF57 Cloning

The ORF57 DNA plasmid was bought from Addgene (pCDNA4.TO-ORF57-2xCSTREP Plasmid #136217). ORF57 was amplified for Gibson Assembly (NEBuilder® HiFi DNA Assembly Master Mix #E2621) using ORF57_F (5’-3’ggatctggagctctcgagaattctcacgcgtCATATGGCCACCatggtacaagcaatgatag) and ORF57_R (5’-3’ CTTGATGATGGCCATGTTATCCTCCTCGCCCTTGCTCACACTAGTGGAACCACCACCACCagaaagtggataaaagaataaaccc). This protein has been optimized for expression in mammalian cells by addition of a SFFV promoter, Kozak sequence, and ATG start codon. The amplified fragments were assembled into a lentiviral packaging plasmid, pHR-SFFV-mCherry, with a C-terminus mCherry tag. mCherry tag was chosen for its monomeric nature and resistance to aggregation, as much of the ORF57 studies performed were tracking pathological aggregation.

### Lentivirus

Plasmids pHR-SFFV-ORF57::mCherry (ORF57::mCherry) and pHR-SFFV-mCherry (mCherry) were produced using the MACHEREY-NAGEL NucleoBond Xtra Maxi kit for transfection-grade plasmid DNA (#740414.50) per the protocol.

HEK-293T cells were transfected with FuGene HD reagent (Promega # E2311) and PSP plasmid, VSV-G plasmid, and OptiMEM media, at a 1:1:3 ratio, respectively. Lentivirus was harvested from the media at 72 hours concentrated by LentiX concentrator (Takara # 631232) per the protocol. The concentrated lentivirus was resuspended in 1/10 of the original media volume using 1x phosphate buffered saline (PBS) with 25 mM HEPES (Sigma-Aldrich # H3375), pH 7.4.

Lentiviral aliquots of ORF57::mCherry and mCherry were diluted 1:10 and cells were infected using a double infection protocol. Infected cells were allowed to grow in the incubator for 48 hours. After 48 hours, cells were passaged then re-infected 24 hours later, cells were incubated for 48 more hours before experiments.

### qPCR

RNA was extracted using Qiagen RNAeasy Mini Prep Plus Kit and eluted in 30 µl RNAse-free water. 1µg of RNA was amplified into cDNA using Applied Biosystems High Transcription cDNA Synthesis Kit (Thermo Fisher # 4368814). cDNA was amplified using SSO Advanced SyBr Green on the QuantFlex 12K. See Table 1 for primers. Ct levels of ORF57 were normalized with GAPDH and fold change was determined. Ct values above 40 were recorded as 40 for calculations.

### Sodium Arsenite Treatment and Time Course

Sodium arsenite (SA) has been shown to be effective *in vitro* to induce the ISR and SG formation by phosphorylation of eIF2α (Arimoto et al., 2008; Sama et al., 2013). Cells were stressed with 300 µM SA in complete media for 90 minutes.

Time course experiment for SA stress, TDP-43ΔNLS cells expressing mCherry or ORF57::mCherry were seeded into 8-chamber slides at ∼80% confluency. Cells were induced with doxycycline for 24 hours, then 300 µM SA stress was added for 90 minutes. The cells were washed in DBPS and allowed to recover in complete media for 15 minutes, 30 minutes, 1 hour, 2 hours, 3.5 hours, and 6.5 hours. Cells were fixed at each time point and ICC performed.

### BCA Assay

Cells were lysed in RIPA buffer containing 50mM Tris-HCl, 150 mM NaCl, 1% NP40, 0.5 mM EDTA, 0.1% sodium deoxycholate, 0.1% SDS. To RIPA buffer, fresh PhosSTOP (Millipore Sigma # 490684500), Proteinase Inhibitor, and 1mM Pefabloc SC (Millipore Sigma # 30827-99-7) was added directly before addition to cells. Cells were lysed by homogenization on ice and then centrifuged at 10,000xg to pellet cell debris. Supernatant was collected and used for Pierce BCA Protein Assay Kit protocol (ThermoFisher #23225) per protocol. Absorbance was then measured using Softmax Plate Reader. Data was analyzed to quantify protein levels of each sample.

### Immunoblot

Cell lysates were thawed on ice. 10 µg of each sample was added to a solution of 1x NuPAGE LDS Sample Buffer (Novex #1249698) and 1x Bolt Sample Reducing Agent (Novex #B009). Samples were boiled for 5 minutes at 95°C. Boiled samples were then loaded into Invitrogen Blot 4 – 12% Bis-Tris Protein Gel (Invitrogen #NW04125BOX) in 1XMOPS solution. Gel was transferred to a nitrocellulose membrane using the iBlot template #1. Once transferred, the membrane was blocked in 5% non-fat milk in 0.05% Tween in Tris-Buffered Saline (TBST), or 5% BSA in TBST (for phosphorylated proteins) for one hour while gently shaking. Membrane was then incubated in primary antibody (Anti-phospho-eIf2α (Rabbit, 1:500 dilution, Cell Signaling Technologies #119A11), Anti-eIF2α (Rabbit, 1:500, Cell Signaling Technologies #9722), Anti-TDP43 (Rabbit, 1:4000, ProteinTech #12892-1-AP), Anti-puromycin (Mouse, 1:1000, EMD Millipore #MABE343), Anti-Actin (Mouse, 1:35000, Millipore Sigma #MAB1501), Anti-phospho-PKR (pThr451, Rabbit, 1:1000, Invitrogen #44-668G), Anti-PKR (Rabbit, 1:4000, ProteinTech #18244-1-AP) diluted in TBST overnight at 4°C while gently shaking. Membrane was washed 3 times in TBST for 10 minutes each. Membrane was then incubated in secondary antibody, Donkey Anti-Rabbit or Donkey Anti-Mouse HRP, diluted 1:10,000 in TBST for 1 hour. Secondary antibody solution was removed and membrane was washed 3 more times in TBST for 10 minutes each. Membranes were incubated in Super Signal West Pico PLUS stable peroxide and luminol/enhancer (Thermo Scientific #34580) prior to imaging. Membranes were imaged using BioRad software. Signal accumulation mode was used. Images chosen for analysis were analyzed on ImageJ software to detect density of bands of interest in each lane.

### SUnSET Assay

To assess changes in global translation a SUnSET assay was performed. In this assay, puromycin replaces amino acids in mRNA being actively transcribed and translation is aborted (73). Cells were co-incubated with 10 µg/mL puromycin and 300 µM SA diluted in complete media for 90 minutes. As a negative control, cells were pre-treated for 30 minutes with 30 µg/mL cycloheximide (CHX) diluted in complete media. After CHX treatment, the pre-treatment was removed and replaced with 10 µg/mL puromycin diluted in complete media for 90 minutes. Following all treatments, cells were lysed in RIPA buffer and processed according to the Western Blot protocol aforementioned with primary antibody against puromycin (Mouse, 1:1000, Sigma Aldrich #MABE343).

### Soluble/Insoluble TDP43 Fractionation

Following BCA, 150 µg of cell lysate were brought up to the same volume of 125 µL RIPA buffer. Lysates were sonicated in water bath for three 30 second intervals for a total of 90 seconds of sonication. Lysates were centrifuged at 100,000xg for 30 minutes. Supernatant was separated as soluble fraction. Pellet was resuspended in 100 µL RIPA buffer. Resuspended pellet was sonicated and centrifuged as previously mentioned. Wash step was pipetted off and the insoluble fraction (pellet) was resuspended in 40 μL of 8 M urea (2 M Thiourea, 4% CHAPS, 30 mM Tris HCl, pH 8.5). Equal volumes of the soluble and insoluble fractions were then prepared and run as semi non-denaturing western blot. Cell lysates were combined with NuPAGE LDS Sample Buffer (Novex #1249698) and run directly on Invitrogen Blot 4 – 12% Bis-Tris Protein Gel (Invitrogen #NW04125BOX) in 1XMOPS solution, Density of the whole lane was determined with ImageJ/Fiji. Primary antibody used was Anti-GFP (Rabbit, 1:2000, ProteinTech #50430-2-AP). Secondary antibody used was Donkey Anti-Rabbit diluted 1:10,000 in TBST.

### Immunocytochemistry

Cells expressing mCherry or ORF57::mCherry were seeded into 8-chamber slides at ∼80% confluency. Cells were fixed in 0.5 mL 4% PFA/PBS. PFA solution was removed and slides were washed in PBS. Cells were permeabilized in 0.5 mL PBS/0.1% Triton X-100 (PBS-T). Cells were then blocked with 10% donkey serum (DS) in PBS for 1 hour. Cells were then incubated in primary antibody (Anti-G3BP1 (Rabbit, 1:300 dilution, Millipore Sigma #07-1801), Anti-eIF3ɳ (Mouse, 1:250 dilution, Santa Cruz #137214), Anti-TDP43 C-terminus (Rabbit, 1:1000, ProteinTech #12892-1-AP), Anti-G3BP1 (Mouse, 1:250, ProteinTech #66486-1-Ig)) diluted in 5% DS in PBS overnight at 4°C. After overnight incubation, the cells were washed with PBS, then incubated in secondary antibody (Donkey Anti-Rabbit Alexa Fluor 647 (1:500), Donkey Anti-Mouse Alexa Fluor 488 (1:500)) diluted in PBS for 1 hour at room temperature. After secondary antibody incubation, cells were washed in PBS. Nuclei were stained with DAPI diluted 1:10,000 in PBS for 5 minutes. DAPI was removed and cells were washed 2 final times in PBS for 10 minutes each. Prolong Gold Antifade mounting media (Invitrogen # P36930) was added before coverslips. Cells were imaged on Keyence microscope. Images were analyzed to find aggregates per cell of stained proteins using Aggrecount v1.1 in ImageJ/Fiji. Mander’s coefficient was calculated by converting images to 16-bit and then using JACoP in ImageJ/Fiji.

### TDP-43 Time Course

TDP-43 NLS cells expressing either mCherry or ORF57::mCherry were seeded in triplicate in an 8-chamber slide. TDP-43 was induced with doxycycline for 24 hours and then the cells were stressed with 300 μm sodium arsenite for 90 minutes. The cells were then rinsed with DPBS and then allowed to recover in complete media for 0 minutes, 15 minutes, 30 minutes, 1 hour, 2 hours, 3.5 hours, or 6.5 hours. The cells were fixed and imaged per the immunocytochemistry protocol above.

### Statistical Analysis

Statistical analysis was performed in GraphPad/Prism v9. Two-way ANOVAs and Bonferroni post hoc tests were used to compare means between conditions when there was a significant interaction. Unpaired t-tests were used for analysis of SUnSET and TDP-43 fractionation immunoblots. P value of 0.05 was used for significance and mean ± standard deviation was reported for ANOVAs. The time course experiment was analyzed using exponential one-phase decay with a least squares regression. The best-fit curves were compared between data sets to find if there was one curve that adequately fit both data sets. Results were visualized in graphs made from GraphPad/Prism.

### Immuno-purification and sample preparation for LC-MS/MS

Cells expressing either mCherry or ORF57::mCherry were plated in four replicates in 6-well plates. Once these cells reached confluency, a subset of these cells were treated with 300 µM of sodium arsenite for 90 minutes to induce cellular stress. Subsequently, these cells were lysed in IP buffer: 10 mM Tris/Cl pH 7.5, 150 mM NaCl, 0.5 mM EDTA, 0.5 % Nonidet™ P40 Substitute, 1% CHAPS and the protein levels were quantified using BCA assay.

Equal amounts of proteins were immunoprecipitated overnight at 4C using the RFP-trap magnetic agarose beads, as recently described by our laboratory (Chromotek, RFP-TRAP Agarose, Cat#rta) (62). The following day, these beads were washed two times with lysis buffer (IP buffer) and they were then washed twice with cold-phosphate buffer saline (pH 7.4). Subsequently, these beads were stored at –80°C for further analysis.

Beads were suspended in 200 µL of ammonium bicarbonate (50 mM) to perform on bead-digestion. The suspended beads were incubated for 30 minutes at room temperature with both 40 mM chloroacetamide and 10 mM TCEP to alkylate and to reduce the proteins, respectively. Then, the reaction was quenched using 20mM DTT. Subsequently, these beads were digested overnight at 37°C with 1 µg of MS-grade trypsin (Thermo Fisher Scientific, Cat#90058). The following day, the digested peptides were collected into new tubes and the trypsin-digestion reaction was quenched with formic acid (final concentration of 1%). These peptides were desalted using Pierce C18 columns (Thermo Fisher Scientific, Cat#89870) following manufacturer’s protocol. Finally, desalted peptides were stored at –80C for further proteomics analysis.

### HPLC-ESI MS/MS and Data Analysis

The LC-MS/MS analysis was performed as previously described (62). Briefly, desalted peptides were re-dissolved in 1% formic acid and fractionated using C18 PepMap pre-column (3µm, 100Å, 75mm x 2cm) hyphenated to a RSLC C18 analytical column (2mm, 100 Å, 75 µm x 50cm). High performance nanoflow liquid chromatography-Orbitrap tandem mass spectrometry (LC-MS/MS) were performed using the Easy nLC 1200 system coupled to Q-Exactive HF-X MS (Thermo Scientific).

Raw data were searched against the Human proteome (the 2018_04 Uniprot release of UP000005640_9606) using MaxQuant software (version 1.6.7.0), with match-between runs activated. All other settings were left at default. The Mass Spectrometry Downstream Analysis Pipeline (MS-DAP, version 1.0.2), which is available at https://github.com/ftwkoopmans/msdap; was used for quality control and differential testing (74). Peptides present in ≥ 75% of sample replicates per contrast and proteins identified by > 2 peptides, were used for differential testing. If the peptides did not match by ≥ 75% of sample replicates between groups, then the analysis was left blank. The Variance Stabilizing Normalization and mode between protein algorithms were used for normalization. Statistical testing was performed with the DEqMS package after rollup to proteins. All obtained raw data, and the complete quality control and differential testing report have been deposited to the ProteomeXchange Consortium via the PRIDE (75,76) (partner repository with the dataset identifier PXD039246. Enriched proteins from the comparisons mCherry vs ORF57 and mCherry-SA vs ORF57-SA, using a FDR-adjusted p-value < 0.05 and Log_2_ fold-change > 0.58 versus the mCherry controls, were cross-referenced with the human TARDBP interactome (BioGRID, accessed on 11.21.2022). Additionally, these enriched proteins were used to build protein networks using Cytoscape Version 3.9.1 and this network was sub-clustered with the ClusterMaker2 Version 2.3.2 plugin. Functional annotation terms were determined using ShinyGO Version 0.76.3 (77).

## ABBREVIATIONS

AD: Alzheimer’s disease
ALS: amyotrophic lateral sclerosis
CHX: cycloheximide
ER: endoplasmic reticulum
FTD: frontotemporal dementia
FTD-TDP: frontotemporal dementia-TDP-43
GCN2: general control nonderepressible 2
HRI: heme-regulated eIF2α kinase
ISR: integrated stress response
PERK: PKR-like endoplasmic reticulum kinase
PKR: protein kinase R
RBP: RNA binding protein
TBST: Tris-Buffered Saline
TDP-43: TAR DNA-binding protein 43kDa

## Acknowledgements

We thank Aquinnah Pharmaceuticals Inc., for use of the ΔNLS-TDP-43 SH SY5Y cell line. B.W. was supported by BrightFocus Foundation (AN2020002) as well as NIH (AG080810, AG050471, AG056318, AG064932, AG072577). A.R-O. was supported by the Rainwater Charitable Foundation.

